# The origin of domestication genes in goats

**DOI:** 10.1101/2020.01.14.905505

**Authors:** Zhuqing Zheng, Xihong Wang, Ming Li, Yunjia Li, Zhirui Yang, Xiaolong Wang, Xiangyu Pan, Mian Gong, Yu Zhang, Yingwei Guo, Yu Wang, Jing Liu, Yudong Cai, Qiuming Chen, Moses Okpeku, Licia Colli, Dawei Cai, Kun Wang, Shisheng Huang, Tad S. Sonstegard, Ali Esmailizadeh, Wenguang Zhang, Tingting Zhang, Yangbin Xu, Naiyi Xu, Yi Yang, Jianlin Han, Lei Chen, Joséphine Lesur, Kevin G. Daly, Daniel G. Bradley, Rasmus Heller, Guojie Zhang, Wen Wang, Yulin Chen, Yu Jiang

## Abstract

Goat domestication was critical for agriculture and civilization, but its underlying genetic changes and selection regimes remain unclear. Here we analyze the genomes of worldwide domestic goats, wild caprid species and historical remains, providing evidence of an ancient introgression event from a West Caucasian tur-like species to the ancestor of domestic goats. One introgressed locus with a strong signature of selection harbors the *MUC6* gene which encodes a gastrointestinally secreted mucin. Experiments revealed that the nearly fixed introgressed haplotype confers enhanced immune resistance to gastrointestinal pathogens. Another locus with a strong signal of selection may be related to behavior. The selected alleles at these two loci emerged in domestic goats at least 7,200 and 8,100 years ago, respectively, and increased to high frequencies concurrent with the expansion of the ubiquitous modern mitochondrial haplogroup A. Tracking these archaeologically cryptic evolutionary transformations provides new insights into the mechanism of animal domestication.

**One Sentence Summary:** Goat domestication mainly focused on immune and neural genes, with adaptive leaps driven by introgression and selection.

## Introduction

Domestication presents an extreme shift of physiological and behavioral stress for free-living animals (*1*) to a high-density and disease-prone anthropogenic environment, especially for herbivores. The goat (*Capra hircus*) was one of the first domesticated livestock species, demonstrating remarkable adaptability and versatility (*2, 3*), and has been intimately associated with human dispersal (*4*). Recent studies have identified candidate targets of selection during goat domestication, including loci related to pigmentation, xenobiotic metabolism and milk production (*5, 6*); however, the evolutionary dynamics of key genes involved in adaptation during the early phase of domestication remains unclear.

Goat domestication is believed to have begun ~11,000 years ago from a mosaic of wild bezoar populations (*Capra aegagrus*) (*2, 6*). However, other wild *Capra* species are widely distributed and many of them can hybridize with domestic goats (*7–9*). Their contribution to the goat domestication process remains unexplored. Introgression of adaptive variants has been widely recognized as a significant evolutionary phenomenon in humans (*10*), sheep (*11*), and cattle (*12*), and it may be particularly effective for increasing fitness without negative pleiotropic effects as demonstrated in other species (*13*). Here, we conducted a comprehensive population genomic survey of modern goat populations distributed across the world, six wild *Capra* species and previously published ancient goat genomes to investigate temporal shifts in key genetic variants under selection during goat domestication.

## Results

### Population structure and origin of domestic goats

We sequenced 101 *Capra* genomes (coverage 3 - 47×, average 12×), including 88 domestic goats collected from three different continents, one bezoar, one Alpine ibex (*Capra ibex*), three Siberian ibexes (*Capra sibirica*), three Markhors (*Capra falconeri*), one West Caucasian tur (*Capra caucasica*) and four Nubian ibex × domestic goat hybrids (*Capra nubiana* × *C. hircus*) (**Fig. 1A, figs. S1 and S2, tables S1 and S2, and text S1**). We also sequenced five ancient goat samples at ~0.04 to 13.44× coverage (**Fig. 1A, figs. S3 to S6, and table S3**), including the earliest known Chinese archaeological samples in Neolithic age (*14*). Together with the publicly available genomic sequences for modern goat and bezoar (*5, 15*), as well as ancient genomes (*6*) (**table S4**), we compiled a worldwide collection of 164 modern domestic goats, 52 ancient goats, 24 modern bezoars and four ancient bezoars.

**Fig. 1.**
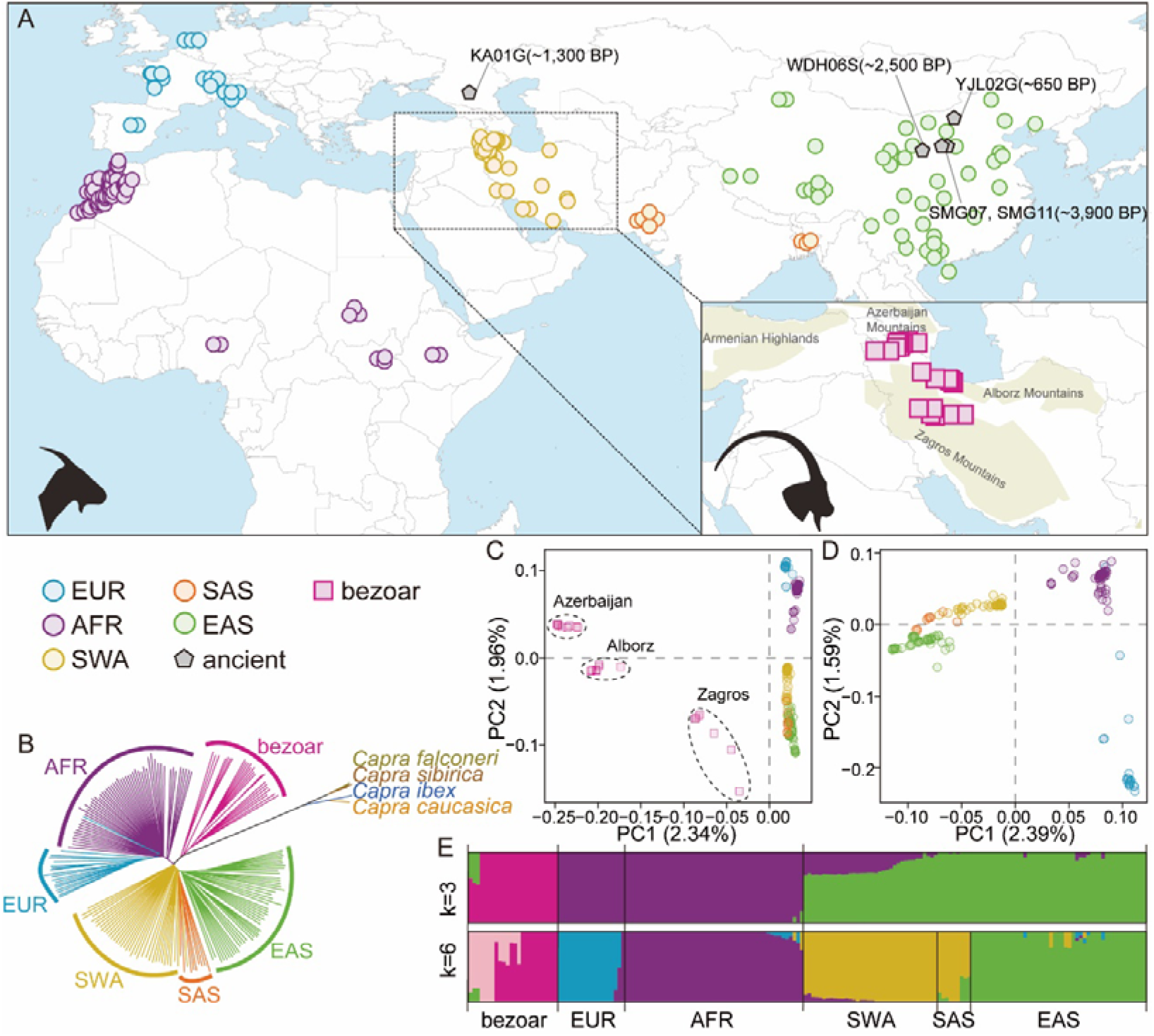
Geographic distribution and genetic affinities of wild and domestic goats used in this study. (**A**) Locations of wild (squares), domestic (circles), and ancient (pentagon) goats for each major geographic group. Domestic goats are colored to mirror their geographic origin. (**B**) A neighbor-joining tree from the genome sequences used in this study. The branches are colored following the same color code used in panel (**A**). (**C** and **D**) Principal component analysis (PCA) of bezoars and domestic goats (**C**) and domestic goats only (**D**). (**E**) ADMIXTURE results for *k* = 3 and *k* = 6, which had the low cross-validation error.

A whole-genome neighbor-joining phylogenetic tree reveals that all domestic goats form a monophyletic sister lineage to bezoars (**Fig. 1B, fig. S7, and table S5**), confirming that modern domestic goats are the descendants of bezoar-like ancestors (*16*). The other four wild *Capra* species (*C. ibex, C. sibirica, C. falconeri* and *C. caucasica*), which are referred to as the ibex-like species (*17*), fall exclusively in another branch divergent from bezoar-goat (**Fig. 1B and fig. S7**). Principal component analysis (PCA) shows that the three bezoar populations, which were collected in the Middle East (**Fig. 1A**) near the domestication center (*2*), are structured and cluster corresponding to their geographic origins (**Fig. 1C**). Within domestic goats, PCA and model-based clustering (*k* = 3) show that Asian goats are genetically distinct from European (EUR) and African (AFR) samples (**Figs. 1D and 1E**). At *k* = 6, Asian goats further split into two geographic subgroups: Southwest Asia-South Asia (SWA-SAS) and East Asia (EAS) (**Fig. 1E and figs. S8 to S10**). This geographic structure is also supported by TreeMix and haplotype-based statistics (**figs. S11 and S12**), in agreement with the scenario that the ancestors of present-day domestic goat populations followed distinct dispersal routes along the east-west axis of Afro-Eurasia (*4*) (**Fig. 2A, figs. S13 and S14, and table S6**).

Our demographic analyses using MSMC, SMC++, and *∂a∂i* indicate that the divergence times amongst Asian, African and European goat populations predated the archaeologically estimated domestication time by a large margin (**figs. S11 to S17, table S7, and text S2**). In addition, the three bezoar populations share significantly different levels of alleles with divergent goat populations according to *D* statistics (**Fig. 2B and fig. S22**). Therefore, the deep coalescence times of different domestic goat populations can be explained by their origin in structured ancestral bezoar populations (*6*), or by post-domestication recruitment from different local bezoar populations.

**Fig. 2.**
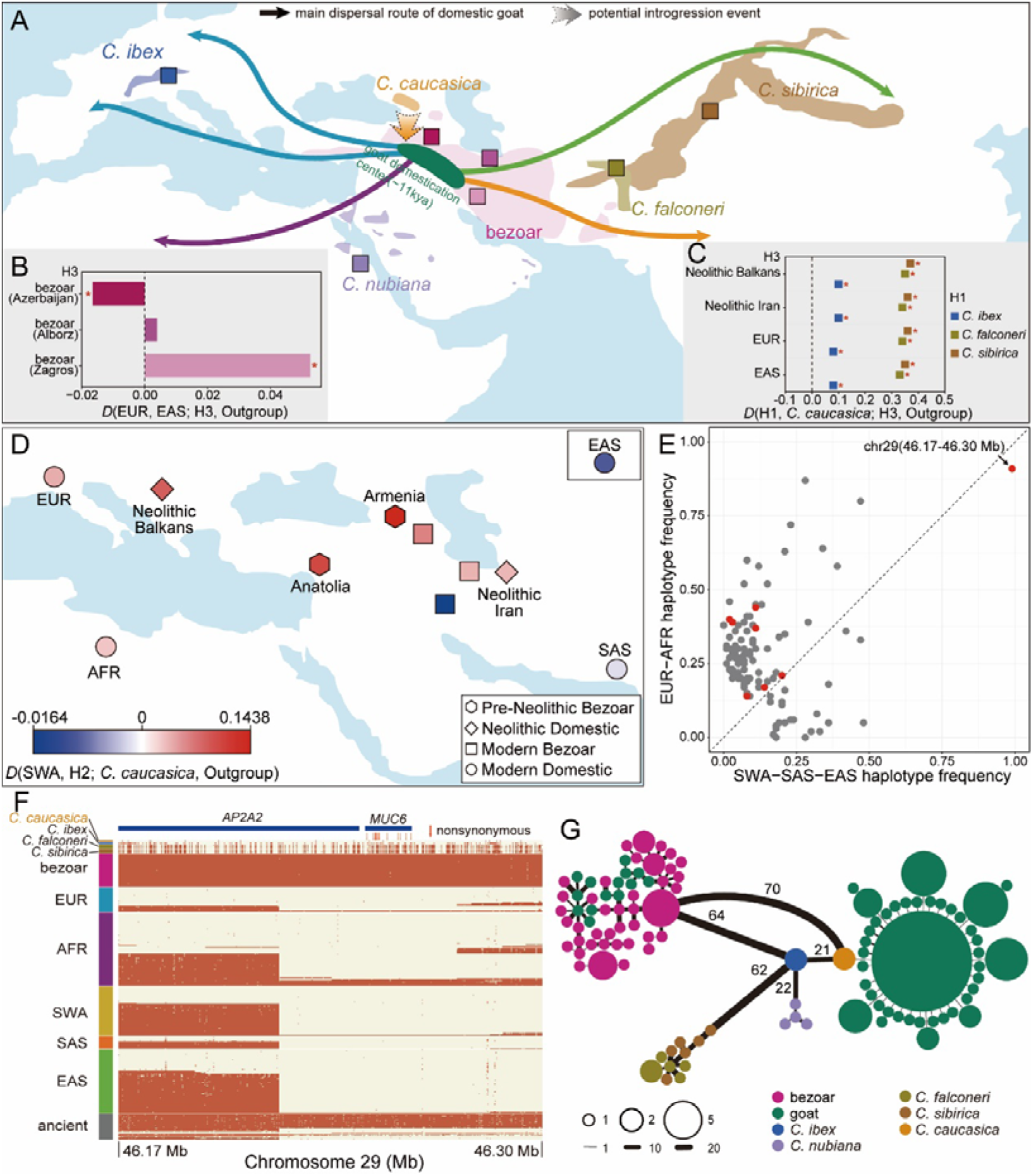
Gene flow during the early stage of goat domestication and diffusion. (**A**) Geographic distribution of the wild *Capra* species and the dispersal routes of domestic goats out of their domestication areas (*3, 7*). (**B**) Allele sharing between domestic goats and bezoars. (**C**) Allele sharing between domestic goats and ibex-like species. Statistically significant results, defined by |Z-scores| ≥ 3, are marked with a red asterisk for panel (**B**) and panel (**C**). (**D**) A heatmap of *D* statistic testing for the differential affinity between SWA (blue) and H2 (red), where H2 represents the individual/population of bezoars and domestic goats located in the map. The East Asian goats are depicted in the box in the upper right corner. Positive *D* (H1, H2; H3, Outgroup) values indicate more allele sharing between H2 and H3 while negative *D* values indicate more allele sharing between H1 and H3. (**E**) A scatter plot of introgressed haplotype frequency in EUR-AFR and Asian goat populations. The red dots represent immune-related loci. (**F**) Haplotype pattern at the *MUC6* locus defined by putatively introgressed variants (yellow, predicted introgressed allele; red, predicted non-introgressed allele). (**G**) Haplotype network from the number of pairwise differences at the *MUC6* non-repeat region. All domestic goats are colored in green, and wild *Capra* species are divided into five subgroups, including the haplotypes from hybrids of Nubian ibex × domestic goat. The radius of the pie chart and the width of edges were log2-transformed.

### Gene flow from ibex-like species to pre-domestication bezoars and modern goats

*D* statistics reveal that all four ibex-like species have different signals of allele sharing with ancient and modern goats, with all goat populations showing the greatest sharing with the West Caucasian tur (**Fig. 2C and table S8**). Notably, all four pre-domesticated bezoars (an Armenian bezoar dating to >47,000 years ago and three Anatolian bezoars dating to ~13,000 years ago) show an excess of allele sharing with the West Caucasian tur (**Fig. 2D**). Further *f*_3_ statistics in the form of *f*_3_ (modern bezoar, West Caucasian tur; target) support gene flow from the West Caucasian tur to ancient Armenian bezoar, but do not suggest additional recent gene flow to modern goats (**table S9**). Together with the fact that three pre-domestication Anatolian bezoars possessed a tur-like mitochondrial haplotype (*6*), this evidence supports a pre-domestication admixture event of bezoar with a tur-like species. This ancient admixture event is further supported by the distribution of West Caucasian tur being sympatric with that of the ancient Armenian bezoar, and closer to the goat domestication center than any other living ibex-like species (**Fig. 2A**).

Although all modern goat genomes suggest some introgression from tur, Neolithic Balkan and modern EUR and AFR goats show more allele sharing with tur compared to Asian goats (**fig. S23**), probably because of higher genetic affinity with the ancient Armenian and Anatolian bezoars (*6*) (**Fig. 2D**). Therefore, pre-domestication introgression from a tur-like species may have contributed alleles to ancient bezoars and thereby early domesticated goats and their derived modern populations.

We identified a total of 112 putative introgressed regions derived from ibex-like species with a haplotype frequency > 0.1 in domestic goats through *D* statistics, Identity-By-State (IBS), Sprime, and maximum likelihood (ML) phylogenetic trees (**Fig. 2E, fig. S24, and Data file S1**). The 112 regions cover 81 protein-coding genes.

A KEGG pathway enrichment analysis for these genes shows that the most significantly enriched category is amoebiasis (hypergeometric test, adjusted *P* < 5.28 × 10^-3^, **table S10**), which is related to parasite invasion and immunosuppression, including four genes (*SERPINB3, SERPINB4, CD1B*, and *COL4A4*). Three additional introgressed genes (*BPI, MAN2A1*, and *CD2AP*) are also involved in immune function (*18–20*). For half of the introgressed regions, the West Caucasian tur shows the highest similarity with the introgressed alleles among the four ibex-like species (**fig. S25 and Data file S1**). Furthermore, the EUR-AFR populations have a higher prevalence of introgressed regions than the Asian populations (*t*-test, *P* = 3.30×10^-10^) (**Fig. 2E**), consistent with the higher genome-wide allele sharing between EUR-AFR and West Caucasian tur.

Notably, one introgressed region on chromosome 29 has a nearly fixed introgressed haplotype (frequency = 95.7%) in the modern worldwide domestic goats (**Figs. 2D and 2F**). This region contains one intact protein-coding gene, *MUC6*, which encodes a gastrointestinally secreted mucin that forms a protective glycoprotein coat involved in host innate immune responses to the invasion of multiple gastrointestinal pathogens (*21–23*). We searched for potential donor populations in our sequenced wild caprid genomes and used coalescent-based inference methods to validate this introgression signal. The haplotype network constructed with the non-repeated region of *MUC6* showed the most common domestic haplotype (*MUC6*^D^) is similar to West Caucasian tur, differing by only one mutation, while being strikingly different from the minor frequency haplotypes of domestic goats and the common haplotype in bezoar (*MUC6*^B^) (**Fig. 2G**). Coalescence time estimates and neutral simulations suggest that such pattern of haplotype differentiation is highly unlikely in the absence of interspecific gene flow (**figs. S26 and S27**). Therefore, the nearly fixed *MUC6*^D^ in goats was most likely introgressed from a lineage close to the West Caucasian tur, in agreement with the genome-wide signal.

### Domestication-mediated selection on immune and neural genes

To identify key selective sweeps in goat domestication, the worldwide domestic goat populations were compared with all 24 bezoars by estimating pairwise genetic differentiation (*F*_ST_), differences in nucleotide diversity (π ln-ratio bezoar/domestic) and cross-population extended haplotype homozygosity (XP-EHH) in 50 kb sliding windows along the genome (**figs. S28 and S29 and table S11**). We defined the windows with significant values (Z test, *P* < 0.005) in all three statistics (*F*_ST_ > 0.195, π ln-ratio > 0.395 and XP-EHH > 2.10) as putative selective sweeps. After merging consecutive outlier windows, 105 unique regions containing 403 protein-coding genes were identified (**Fig. 3A and Data file S2**). 18 of these regions have been identified in previous domestication scans, including those associated to phenotypic effects related to immunity, neural pathways/processes, pigmentation, and productivity traits associated with milk composition and hair characteristics (**Data file S2**). The limited overlap with previous studies is mainly due to differences in sample sets, selection scan methodology and different versions of the reference genome (**text S2**).

**Fig. 3.**
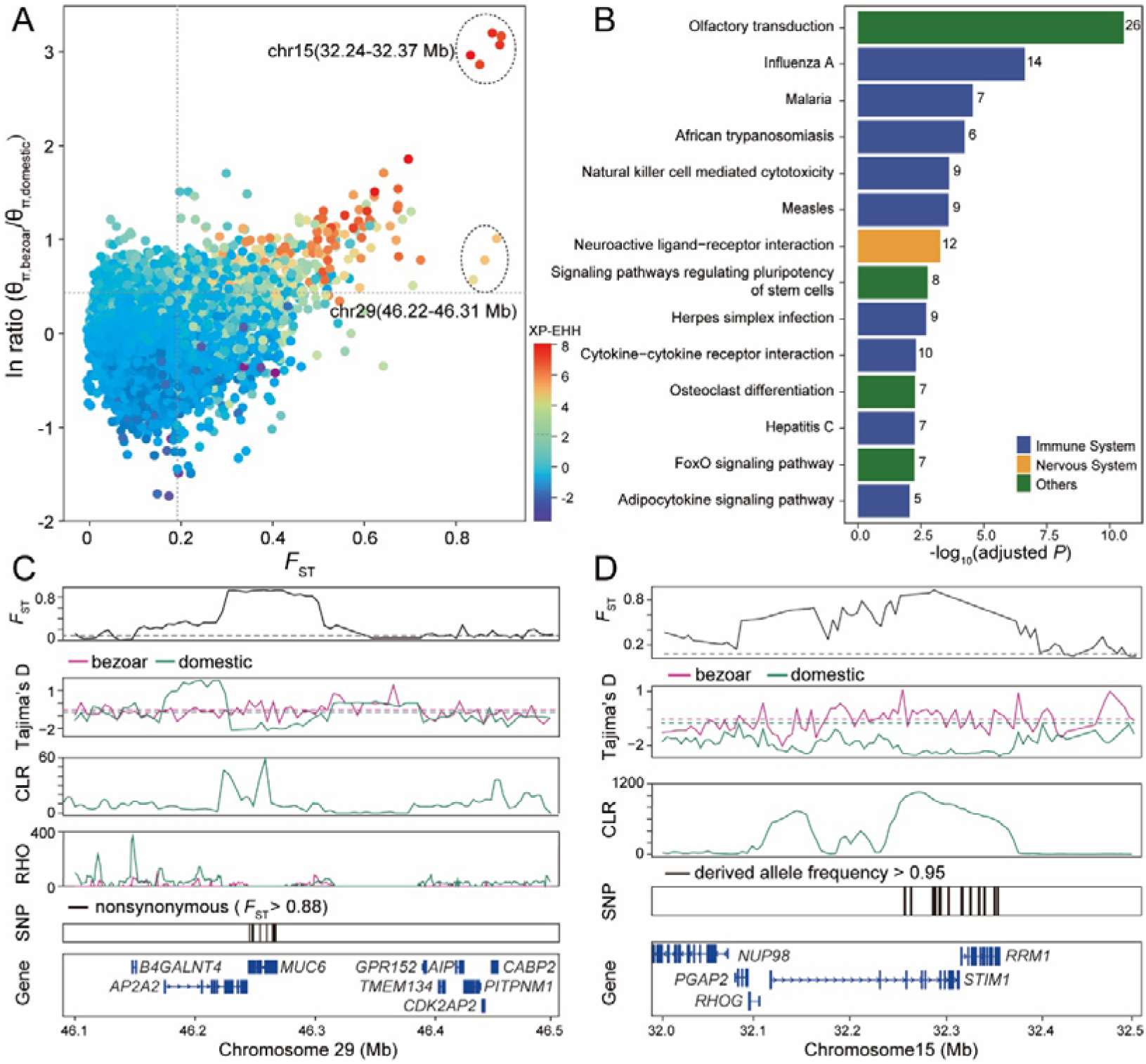
Genomic regions with selection signals in domestic goats. (**A**) Distribution of the pairwise fixation index (*F*_ST_) (x-axis), π ln ratio (y-axis) and value of XP-EHH (color) between bezoars and domestic goats. The dashed vertical and horizontal lines indicate the significance threshold (corresponding to Z test *P* < 0.005, where *F*_ST_ > 0.195, π ln ratio > 0.395, and XP-EHH > 2.1) used for extracting outliers. (**B**) KEGG pathways identified as significantly overrepresented (hypergeometric test, adjusted *P* < 0.01). (**C** and **D**) Selective sweep and distribution of the recombination (RHO) on chromosome 29 (chr29:46.22-46.31 Mb) (**C**) and selective sweep on chromosome 15 (chr15:32.24-32.37 Mb) (**D**). The putative sweep region is additionally validated by *F*_ST_, Tajima’s D and composite likelihood ratio (CLR) test. Horizontal dashed lines represent the whole-genome mean for the corresponding parameters. Gene annotations in the sweep region, nonsynonymous SNPs with *F*_ST_ > 0.88 (**C**), and SNPs nearly fixed for derived alleles in domestic goats (**D**) are indicated at the bottom.

KEGG analysis showed that nine of the 14 significantly enriched pathways (hypergeometric test, adjusted *P* < 0.01) are immune-related (hypergeometric test, adjusted *P* = 5×10^-3^ to 2.35×10^-7^) (**Fig. 3B and table S12**), including influenza A, malaria, African trypanosomiasis, Natural killer cell-mediated cytotoxicity, Measles, Herpes simplex infection, Cytokine-cytokine receptor interaction, Hepatitis C, and Adipocytokine signaling pathway. We surveyed the literature and identified 40 additional genes with immune function (**table S13**), obtaining a total of 41 regions that contain 77 immune-related genes. Among these, *F*_ST_ is particularly high at *MUC6* region and we detected 16 missense mutations showing *F*_ST_ > 0.88 in this gene (windowed *F*_ST_ = 0.89, π ln-ratio = 1.01, XP-EHH = 5.25. **Fig. 3C, fig. S30, and table S14**). This region contains 228 SNPs nearly fixed for the derived allele in modern goats (frequency > 95%, absent in bezoars) accounting for 93.8% of those in the whole genome (a total of 243, **Data file S3**), illustrating the singular nature of this selection signal.

The selection enrichment analysis furthermore pinpoints 12 genes involved in Neuroactive ligand-receptor interaction (hypergeometric test, adjusted *P* = 3.70×10^-5^). We observed an additional 37 genes linked to other functions in the nervous system that were under selection during goat domestication (**table S15**). One genomic region located on chromosome 15 shows the strongest combined signal of selection (*F*_ST_ = 0.90, π ln-ratio = 3.20, and XP-EHH = 8.09) (**Fig. 3A**). Evidence for large negative Tajima’s D values, high composite likelihood ratio (CLR) scores, and extensive haplotype sharing in domestic goats all suggest strong positive selection at this locus (**Fig. 3D and fig. S31**). This locus contains 15 SNPs almost fixed for the derived allele in domestic goats and harbors two protein-coding genes, *STIM1* and *RRM1* (**Data file S3**). Functionally, *STIM1* is an endoplasmic reticulum calcium sensor involved in regulating Ca^2+^ and metabotropic glutamate receptor signaling in the neural system (*24, 25*). *RRM1* encodes ribonucleoside-diphosphate reductase large subunit, an enzyme essential for the production of deoxyribonucleotides, and influences sensitivity to valproic acid induced neural tube defects in mice (*26*). Hence, we hypothesize that this strongly selected locus could be an example of behavioral adaptions in the early phase of animal domestication (*27*).

### Introgressed *MUC6* plays a role in pathogen resistance

The *MUC6* region constitutes the only intersection between the selection and introgression scans. In sheep and cattle, *MUC6* is associated with gastrointestinal parasite resistance (*28–31*). The results of transcriptome sequencing, qPCR, and immunohistochemistry revealed that *MUC6* is specifically expressed in the abomasum and duodenum of goat (**Fig. 4A and figs. S32 and S33**). Structurally, MUC6 has a long variable number of tandem repeats (VNTRs) rich in Pro, Thr, and Ser residues, which can influence the covalent attachment of O-glycans (*32*). We therefore used PacBio SMRT transcriptome sequencing to investigate the difference between the *MUC6* and *MUC6* at the transcriptional level. Besides a number of small indels, an 82 amino acid (aa) deletion containing three copies of the VNTR after the 2,789th aa was uniquely identified in *MUC6*^D^ compared to *MUC6*^B^ (**figs. S34 and S35**). The number of VNTRs in these two haplotypes may influence the function of *MUC6*, which is a key to the generation of gastrointestinal mucous gel (*22, 33*). To examine the potential difference of pathogen resistance of the *MUC6* and *MUC6* variants, an epidemiological survey was carried out in a polymorphic goat population. By evaluating fecal egg counts (FEC) for gastrointestinal nematodes in 143 *MUC6*^D^/*MUC6*^D^, 111 *MUC6*^D^/*MUC6*^B^, and 14 *MUC6*^B^/*MUC6*^B^ ewes at 13 months of age (**fig. S36**), we found that the goats carrying *MUC6*^D^/*MUC6*^D^ exhibited lower FEC than those carrying the other two genotypes (**Fig. 4B and Data file S4**), implying that the goats carrying *MUC6*^D^ might be more resistant to gastrointestinal nematodes. These results support that the introgressed *MUC6*^D^ in domestic goats is most likely under selection due to its advantage in the host innate immune response towards potential gastrointestinal pathogens (*22*).

**Fig. 4.**
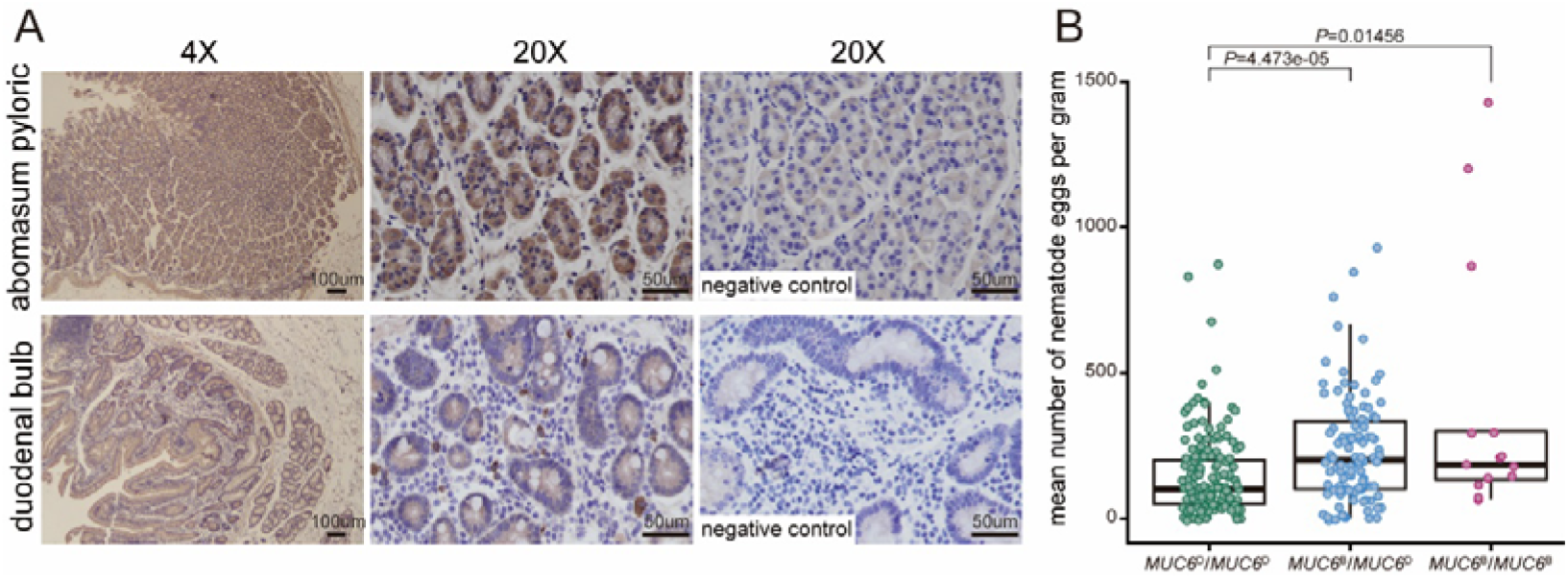
The association of *MUC6* with gastrointestinal nematodes resistance. (**A**) Immunohistochemistry for MUC6 in abomasum pyloric and duodenal bulb of a goat. Photomicrographs at 4X, 20X are shown on the left and in the middle. The negative controls (20X) are shown on the right. (**B**) The statistical association between *MUC6* genotype and the fecal egg counts for gastrointestinal nematodes. Wilcoxon’s rank-sum test was employed to compute the *P* values.

### The origin and diffusion of domestic *STIM1-RRM1* and *MUC6* alleles

An in-depth genetic survey of ancient genomes throughout the Near East revealed that two ancient Balkan goats dating to ~8,100 years ago carried *STIM1-RRM1*^D^ (**fig. S37 and Data file S5**). *MUC6*^D^ appeared later at ~7,200 years ago in Southwest Iran (**fig. S38 and Data file S6**), a period characterized by an increased density of settlements in the Fertile Crescent (*34*). Herding goats at higher densities in a crowded and disease-prone anthropogenic environment would likely have exerted an increased selective pressure for livestock pathogen resistance (*35*). Interestingly, the first detected ancient animals possessing *STIM1-RRM1*^D^ or *MUC6*^D^ both carry mitochondrial haplogroup A, although this mitochondrial haplogroup had a low frequency and narrow distribution before 7,500 years ago (*6*). By ~6,500 years ago, the frequency of the *STIM1-RRM1*^D^ and *MUC6*^D^ increased to ~60% in the Near East (**Fig. 5A**) and spread to China ~3,900 years ago (**Fig. 5B**), concomitant with the consolidation and diffusion of livestock-based economies throughout Eurasia (*36, 37*). The expansion of these two selected loci was also contemporaneous with the overwhelming spread of mitochondrial haplogroup A. In contrast, the frequencies of the two major Y-chromosome haplogroups remained relatively unchanged over time (**Fig. 5B, figs. S38 to S40, and Data file S7**).

**Fig. 5.**
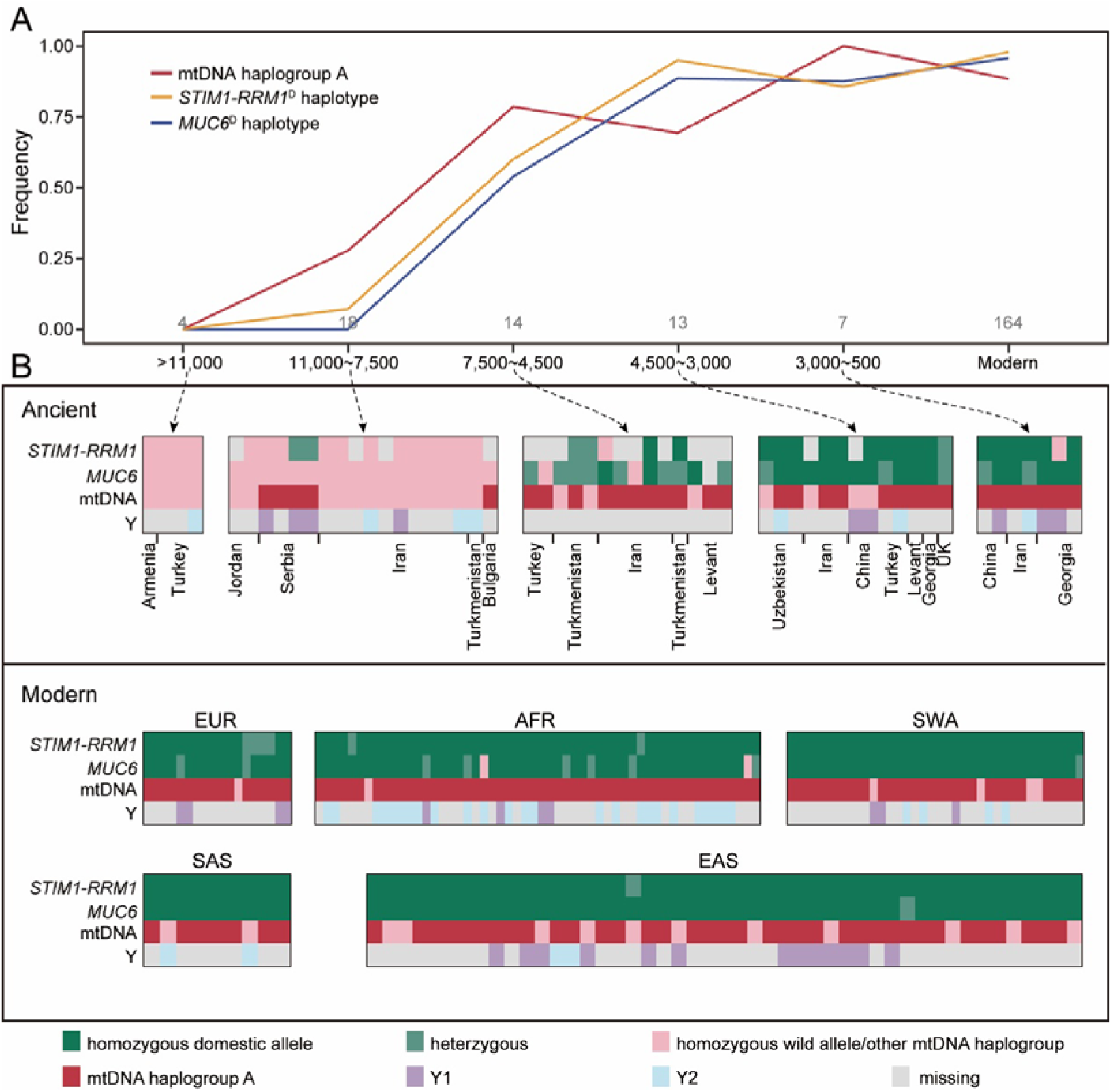
The emergence and diffusion of domestic *STIM1-RRM1* and *MUC6* haplotypes are concurrent with the spread of mtDNA haplogroup A. (**A**) The temporal changes in the frequency of the *STIM1-RRM1D MUC6D* and mtDNA haplogroup A from pre-domestication bezoars to modern domestic goats. The dates are expressed as cal. BP. (**B**) Genotypes of *STIM1-RRM1* and *MUC6*, and mtDNA and Y-chromosome haplogroups. The presences in homozygous or heterozygous states are shown in green and light green, respectively. The absence of the domestic allele is depicted in light pink. The light grey color symbolizes missing information.

## Discussion

The present study generated genomic data from a substantial number of domestic and wild *Capra* species to characterize adaptive introgression and genetic changes during goat domestication. Collectively, both the selective sweep enrichment analyses and the two outstanding selection signals found in *MUC6* and *STIM1-RRM1* in domestic goats are consistent with the hypothesis that genes related to pathogen resistance and behavior have been particularity targeted during goat domestication.

Numerous studies have shown that adaptive introgression can provide beneficial alleles that allow the recipient populations to adapt to new environments (*12, 38–40*). In humans, immune-related loci acquired substantially advantageous alleles by admixture with archaic humans who were probably well adapted to local environments and pathogens (*41–43*). In our study, a number of lines of evidence support that gene flow between West Caucasian tur and goats occurred prior to the onset of domestication. The West Caucasian tur is distributed around the Black Sea coast (**Fig. 2A**), geographically close to the goat domestication center. West Caucasian tur inhabit a humid, subtropical environment where the expected pathogen exposure and parasite load may be considerably higher than in more arid regions of southwest Asia. The rigorous identification of introgression segments in domestic goats shows that the nearly fixed *MUC6*^D^ was introgressed from a West Caucasian tur-like species. These results indicate that the advantageous *MUC6*^D^ might enhance innate immune surveillance and reactivity against potential gastrointestinal pathogens under a grazing agro-ecological niche (*44*) with novel infectious disease pressures.

Our data set including several ancient goat samples allowed us to tentatively track the emergence and spread of the advantageous variants in these two important domestication loci (**Fig. 5**). The first occurrences of the *STIM1-RRM1*^D^ and *MUC6*^D^ locus coincided with the otherwise rare mtDNA haplogroup A, and the three increased in frequency roughly concurrently (**Fig. 5A**), becoming nearly fixed in modern goat populations. These results suggest the global diffusion of variations central to the domestication process was possibly mediated by a subset of female goats carrying the mtDNA haplogroup A. Despite this, modern goat populations remain clearly differentiated from each other. The most likely explanation we can think of is that even Neolithic goat husbandry entailed some kind of breeding strategy under which immigrant matrilines containing globally advantageous alleles were carefully backcrossed with local populations, which probably carried local genetic adaptations, rather than simply replacing local populations.

Overall, our results provide evidence for adaptive introgression and the genetic basis of selected traits during the domestication of goats. We highlight that livestock domestication is a dynamic evolutionary process, with adaptive leaps driven by introgression and selection. Our results indicate that domestication may have focused mainly on neural traits and changes in pathogen resistance, which helped managed herds to adapt to an anthropogenic environment. Our work provides a new insight into goat domestication research and the application of genomics to practical breeding for other domesticated species.

## Materials and Methods

### Sample collection

We sampled 106 animals including 88 domestic goats (*Capra hircus*), one bezoar (*Capra aegagrus*), three Markhors (*Capra falconeri*), three Siberian ibexes (*Capra sibirica*), one Alpine ibex (*Capra ibex*), one West Caucasian tur (*Capra caucasica*), five ancient goats spanning from the Neolithic to the Middle Ages, and four Nubian ibex hybrids (*Capra nubiana* × *C. hircus*) for evolutionary analyses. All procedures involving sample collection were approved by the Northwest A&F University Animal Care Committee (Permit number: NWAFAC1019), making all efforts to minimize invasiveness. The *C. caucasica* (tooth) sample was obtained from a zoo specimen collected by the Muséum national d’histoire naturelle (MNHN, sample ID MNHN ZM AC 1982-1092) in the year 1982. In addition, we also obtained whole-genome data and their geographical coordinates of the sampling sites from the Nextgen project (http://projects.ensembl.org/nextgen) (*5, 15*). Details of the samples used in this study are given in **tables S1 to S3**.

### DNA extraction and sequencing

For modern samples, genomic DNA was extracted from whole blood using the standard phenol-chloroform method and checked for quality and quantity on the Qubit 2.0 fluorometer (Invitrogen). Library construction for resequencing was performed with 1-3 μg of genomic DNA using standard library preparation protocols and insert sizes from 300 to 500 bp. All libraries were sequenced on either the Illumina HiSeq 2500 or X Ten platforms.

For the historical sample (*C. caucasica*), tooth pulverization, DNA extraction and Illumina sequencing library preparation were performed in Trinity College Dublin, Ireland as described by ref. (*6*) and ref. (*45*). The double-stranded DNA sequencing library was constructed without UDG treatment. The sample library was then sequenced using a HiSeq 2500 platform (Single end, 1×101 bp).

### Read alignment and variant calling

Filtered reads from all individuals were aligned to the latest goat reference genome (GCF_001704415.1) (*46*) by the Burrows-Wheeler Aligner (BWA) (*47, 48*). To obtain high-quality SNPs, SNP calling and genotyping were carried out at two separate stages. Variants were discovered on a population-scale (without outgroup argali *Ovis ammon*) using SAMtools model (*49*) implemented in the analysis of next generation sequencing data (ANGSD) (*50*) with “-only_proper_pairs 1 -uniqueOnly 1 -remove_bads 1 -minQ 20 -minMapQ 30 -C 50 -doMaf 1 -doMajorMinor 1 -GL 1 -setMinDepth [Total_read_depth/3] -setMaxDepth [Total_read_depth*3] -doCounts 1 -dosnpstat 1 -SNP_pval 1”. The sites that could be called in at least 90% of the samples and had a strand bias score below 90% were retained. A *P*-value threshold of 1 × 10^-6^ and minor allele frequency value >1/2*n* were used as SNP discovery criteria, where *n* was the sample size. We also kept the sites at which the sampled individuals were homozygous for the alternative alleles. To get the hard-called genotypes for the candidate variants in all *Capra* species and outgroups, we called genotypes according to the GATK-3.7.0 (*51*) best practice workflows. Finally, we only kept biallelic markers as well as markers with no more than 10% missing data using VCFtools v0.1.15 (*52*) with “--min-alleles 2 --max-alleles 2 --max-missing 0.9”. Then, all sample sets of filtered variant calls were used for imputation and phasing using BEAGLE v4.1 (*53, 54*).

To retrieve the nucleotides that were accessible to variant discovery (used in the demographic estimation), hard filtering of the invariant sites was carried out with thresholds based on sequencing coverage (1/3 to 3-fold of corresponding mean depth) and missing data rate (no more than 10%) following the same strategies adopted for variant sites.

### Population structure and phylogenetic analysis

Neighbor-joining (NJ) tree was constructed for the whole-genome SNP set using MEGA v6.0 (*55*) based on pairwise genetic distances matrix calculated by plink v1.9 (*56*). We also made use of the 5,043,096 fourfold degenerate sites to build a maximum likelihood (ML) tree using RAxML v8.2.9 (*57*). PCA was performed using smartpca, part of the EIGENSOFT v6.1 (*58*). A Tracy-Widom test was used to determine the significance level of the eigenvectors. ADMIXTURE (*59*) was used to perform an unsupervised clustering analysis. We increased the number of predefined genetic clusters from *k* = 2 to *k* = 7 and ran the analysis 20 times for each *k*. We further compared the individual-level haplotype similarity using fineSTRUCTURE (*60*). TreeMix v1.12 (*61*) was also used to infer a population-level phylogeny using the ML approach.

### Demographic reconstruction

To infer patterns of effective population size and population separations over historical time, we used MSMC2 (*62*), using the statistically phased autosomal genotypes. We also used the SMC++ (*63*), which does not rely on haplotype phase information. We estimated a mutation rate of 4.32×10^-9^ per site per generation based on the whole genome alignment between goat and sheep (*Ovis aries*) and a generation time of 2 years. Additionally, we calculated the likelihood of the observed site frequency spectrum (SFS) conditioned on several demographic models using *∂a∂i* (*64*). For the best model, nonparametric bootstrapping (100 replicates) was performed to determine the variance of each parameter.

### Gene flow analysis

We computed *f*_3_-statistics (*65*) and *D*-statistics (*66*) to formally test hypotheses of gene flow. To identify regions introgressed into modern domestic goats from ibex-related species, we calculated *D*-statistics using non-overlapping 20 kb sliding windows across the genome. Windows within top 5% of the *D*-statistics were considered as outliers and were further examined. An introgressed segment should have low sequence divergence from the putative donor species whereas sequence divergence between the donor species and bezoar should be higher. Therefore, we also calculated the Identity-By-State (IBS) distance matrix of the outlier regions using ANGSD (*50*). Differences in sequence divergence were determined by a *t*-test-based approach using a *P*-value of 0.01 as cutoff. To further investigate whether introgression best explained the outlier regions, we used Sprime (*67*) to infer the putative introgressed alleles in the significantly diverged regions using 24 modern bezoars as outgroup. Although bezoars may contain a small amount of introgressed variants from ibex-related species, only the frequent (i.e., introgressed allele frequency > 0.1) introgression from other wild *Capra* species to domestic goats were assessed here. Alleles with frequency > 1/48 in the 24 bezoars were excluded from further analysis. We assumed a score threshold of 150,000 and a mutation rate of 4.32 × 10^-9^ per site per generation to determine an introgressed segment (*67*). The region boundaries were defined by the first and last introgressed alleles. To determine putative introgression haplotypes, we required that over 50% of introgressed alleles could be found in introgressed haplotypes. Genomic regions containing at least 100 putative introgressed variants in domestic goats were further examined by constructing ML trees and by searching for domestic goats located within the ibex-related clade. For each region, we also reported a match if the genotype of ibex like species included the putative introgressed allele and a mismatch otherwise. The match rate was calculated as the number of matches divided by the number of compared sites (matches and mismatches).

### Selective sweep analysis

We screened for sweeps selected during goat domestication by parsing specific 50-kb windows that showed low diversity in domestic goats and had high divergence (a high fixation index, *F*_ST_, and haplotype differences, XP-EHH) between domestic goats and wild goats. After calculating all tests, the windows with *P* values less than 0.005 (Z test) were considered significant signals. Candidate genes under selection were defined as those overlapped by sweep regions or within 20-kb of the signals. Two additional statistics, the Tajima’s D and composite likelihood ratio (CLR) test were applied to confirm the top signals.

### Functional enrichment analyses

We characterized the most relevant functions of the protein-coding genes overlapping with the selective sweeps and introgressed regions by searching for overrepresented Kyoto Encyclopedia of Genes and Genomes (KEGG) pathways. Goat protein sequences were used to conduct functional enrichment tests on the target genes using KOBAS 3.0 server (*68*). The *P*-value was calculated using a hypergeometric distribution. FDR correction was performed to adjust for multiple testing. Pathways with an FDR-corrected *P*-value < 0.01 were considered statistically significantly enriched.

### Genotyping loci underlying domestication in ancient genomes

To investigate the genotypes of the *STIM1-RRM1* and *MUC6* locus in ancient samples, we used a total of 243 SNPs (15 SNPs within *STIM1-RRM1* locus and 228 SNPs within *MUC6* locus, respectively), which showed derived allele frequency > 0.95 in 164 modern domestic goats (defined as domestic haplotypes: *STIM1-RRM1*^D^ and *MUC6*^D^) and was absent in 24 modern bezoars and four ancient bezoars (Hovk1, Direkli1-2, Direkli6 and Direkli5) (defined as bezoar haplotypes: *STIM1-RRM1*^B^ and *MUC6*^B^). Due to the low coverage for some ancient genomes, we used allele reads count at SNP positions to determine the genotypes of ancient goats at *STIM1-RRM1* locus and *MUC6* locus. For each ancient goat and SNP, we determined the numbers of reads corresponding to the ancestral and derived allele using GATK UnifiedGenotyper (--min_base_quality_score 15 --output_mode EMIT_ALL_SITES --standard_min_confidence_threshold_for_calling 20) (*51*). For each locus in each ancient goat, we calculated the proportion of the summed derived allele reads count versus total allele reads count. If the proportion was lower than 10%, the genotype of ancient goat was considered to be homozygous for ancestral allele (bezoar-like). The genotype was set to heterozygous if the proportion was between 10% and 90%. If the proportion was more than 90%, it was classified as homozygous for derived allele (domestic-like). We set the genotype to be missing when the ancient goats only had one single informative read (mapping to the predefined SNPs) mapped to the locus.

### Expression and epidemiologic survey related to *MUC6* locus

#### Quantitative real-time PCR analysis

Total RNA was extracted using goat tissues from gastrointestinal tract (including abomasum, duodenum, jejunum, ileum, and caecum) using Eastep Super Total RNA Extraction Kit (Promega, Shanghai, China). cDNA was generated via PrimerScriptTM RT Reagent Kit with genomic DNA Eraser (Perfect Real Time) (TaKaRa, Beijing, China). Quantitative real-time PCR analysis (q-PCR) was performed with FastStart Universal SYBR Green Master (Roche, Shanghai, China) on a Bio-Rad instrument. *MUC6* primers (Forward primer: 5’-CAGCCAGGACAAAATCATGA-3’, Reverse primer: 5’-CTCTGGTCTGGCCTCTGAAC-3’) were designed using NCBI Primer-BLAST (http://www.ncbi.nlm.nih.gov/tools/primer-blast/). *GAPDH* (Forward primer: 5’-ACACCCTCAAGATTGTCAGC-3’, Reverse primer: 5’-ATAAGTCCCTCCACGATGC-3’) was used as internal reference. Gene expression results were calculated using the delta-delta cycle threshold (2^-ΔΔCt^) method.

#### Immunohistochemistry analysis

For histological analysis, dissected goat tissues (abomasum pyloric and duodenal bulb) were fixed in 4% paraformaldehyde, and embedded in paraffin for sectioning (5 μm thick). The tissues sections were deparaffinized and rehydration followed by antigen retrieval using a citrate-buffered solution in a microwave at 100□ for 15 min. After cooling down to room temperature, quenching of endogenous peroxidase and protein block, the sections were treated with a primary antibody (Cat No. D161001, Sangon Biotech, Shanghai, China) overnight at 4□. Negative control sections were obtained by incubating with the primary antibody diluted buffer. Subsequently, the secondary antibody (Cat No. D110058, Sangon Biotech, Shanghai, China) was used to detect primary antibody. Specific protein immunoreactivity was visualized with the substrate chromogen 3,3’-diaminobenzidine (DAB). Finally, the slides were rinsed in water, counterstained with haematoxylin and mounted with coverslips.

#### Full-length transcript sequencing

To determine the sequence differences between the domestic (*MUC6*^D^) and bezoar (*MUC6*^B^) *MUC6* haplotypes, total RNA was isolated from the abomasum pyloric with high expression of *MUC6* from two heterozygous goats and sequenced by PacBio Sequel. Pacbio SMRTbell libraries were sequenced on two separate SMRT cells (Annoroad Gene Technology Co., Ltd., Beijing, China). A total of 23,338,976 subreads were generated with a mean accuracy of 80% and an average length of 1755 nt. High-quality Circular Consensus Sequences (CCS) were obtained using the Iso-Seq 3 application in the PacBio SMRT Analysis v6.0.0 https://github.com/PacificBiosciences/IsoSeq3), with parameters “--noPolish -- minPasses 1”. Finally, we got a total of 960,271 CCS. These CCS were aligned to the latest goat reference genome using Minimap2 (*69*) and we picked out 251 CCS that mapped to the *MUC6* mRNA sequence. We used the CCS that belonged to the bezoar haplotype to manually assemble the mRNA sequence of the *MUC6*^B^. Then, we realigned the *MUC6*^B^ mRNA sequence to *MUC6*^D^ mRNA sequence (XM_018042766.1) by MEGA6 (*55*) and checked the indels carefully. The abundance of tandem repeats in *MUC6* (XP_017898255.1) was examined by using the rapid automatic detection and alignment of repeat finding program (https://www.ebi.ac.uk/Tools/pfa/radar/) (*70*). Three major types of repeat units with distinct sequence features were observed. We detected a 246 bp insertion in the *MUC6*^B^ mRNA sequence located at the 32^nd^ exon, containing three copies of type III units between the 2789th amino acid and the 2871th amino acid. We further validated this insertion by aligning short transcriptome reads from homozygous and heterozygous to the *MUC6*^B^ and *MUC6*^D^ mRNA sequences, respectively.

#### Test population

To further test the genotype-phenotype associations, we first examined the genotypes at the *MUC6* locus using blood samples from breeding rams from a company located in Inner Mongolia, China. This company runs more than 9,000 cashmere goats, mainly for superfine cashmere (fiber diameter < 14 μm). The animals are dewormed routinely with Oxfendazole, Ivermectin, Avermectin, and Levamisole three times annually. Of the 20 tested breeding rams, we identified two animals with *MUC6*^D^/*MUC6*^B^ genotype. We sampled their offspring based on breeding records. Blood samples of these 495 animals were collected in January, 2019.

#### Genotypic and phenotypic analysis

DNA was extracted from the blood samples. The *MUC6* locus was genotyped by PCR (Forward primer: 5’-CAGCACTATCTCCCATACATC-3’, Reverse primer: 5’-GTGGAGCTGAGCTGACACTT-3’) and Sanger sequencing. The frequency of *MUC6*^B^ was 17.3% (n = 338 *MUC6*^D^/*MUC6*^D^, n = 143 *MUC6*^D^/*MUC6*^B^ n = 14 *MUC6*^B^/*MUC6*^B^).

After genotyping, we collected fresh fecal samples from a subset of 268 animals from the rectum in April, 2019, a time when gastrointestinal nematodes numbers were anticipated to be elevated. The gastrointestinal parasitic infestations were examined using the McMaster’s technique as described in ref. (*71*). In brief, two grams of fecal samples were transferred into a container and 60 ml of saturated sodium chloride were added. The suspension was thoroughly stirred with a glass stick and poured through a strainer (80-mesh screen) into the new container. The container with the suspension was closed tightly and carefully inverted several times. Then, the suspension was taken up from the container to fill the glass plate with two McMaster counting chambers. The size of the counting chamber was 10 × 10 × 1.5 mm. The nematodes eggs were then counted under microscopic observation at 100× magnification. Observed nematodes in the feces included the common abomasal and intestinal goat nematodes: *Hemonchus contortus* and *Nematodirus sp*. (**fig. S36**).

The number of nematodes eggs per gram (EPG) was calculated according to the following formula: EPG = (*n*1 + *n*2)/2 ÷ 0.15 × 60 ÷ 2, where (*n*1 + *n*2)/2 is the average number of eggs per chamber, 0.15 is the effective volume (ml) for counting chamber, 60 is the total volume (ml) of suspension, 2 is the weight (g) of faeces examined. EPG was measured on three replicates of each fecal sample and the average of the three replicates was used for analysis.

#### Data analysis

The average fecal nematodes egg counts were not normally distributed, therefore we used Wilcoxon’s rank sum test to test the null hypothesis that there was no association between genotype and phenotype.

## Acknowledgments

We thank members of the Nextgen project for sharing their data. We thank Lobna Mohammed for providing Nubian ibex hybrid blood samples. We thank High-Performance Computing (HPC) of NWAFU for providing computing resources.

## Funding

This project was supported by grants from the National Natural Science Foundation of China (31822052 and 31572381 to Y.J., 31572369 to Y.Chen), the Talents Team Construction Fund of Northwestern Polytechnical University (NWPU) and Strategic Priority Research Program of CAS (XDB13000000) to W.W. and the Villum Foundation (VKR 023447) to R.H.. The Tang scholar at Northwest A&F University (NWAFU) to X.-L.W..

## Author contribution

Y.J. and Y.Chen conceived the project and designed the research. Z.Z., M.L., and X.-H.W., performed the majority of the analysis with contributions from Z.Y., Y.L., Y.Cai, Q.C., J.Liu, K.W., X.P., Y.W., S.H., M.G., T.Z., Y.Z., Y.G., Y.X., and Y.Y.. X.-L.W., W.Zhang, J.H., L.Chen, A.E., and M.O. prepared the modern DNA samples. J.Lesur provided the historical sample of West Caucasian tur. D.C. prepared the ancient samples. X.-H.W., Z.Z., Y.J., and M.L. drafted the manuscripts with input from all authors, and Y.J., W.W., R.H., G.Z, K.D., D.B., L.Colli, and T.S.S. revised the manuscript.

## Competing interests

The authors declare that they have no competing interests.

## Data and materials availability

Individual genome sequence data are available at the Sequence Read Archive (https://www.ncbi.nlm.nih.gov/sra) under accession code PRJNA387635 and PRJNA361447.

## Supplementary Materials

Supplementary Materials and Methods

Supplementary Text

Figs. S1 to S40

Tables S1 to S15

Legends for Data file S1 to S7

